# CLIP-Seq and massively parallel functional analysis of the CELF6 RNA binding protein reveals a role in destabilizing synaptic gene mRNAs through interaction with 3’UTR elements *in vivo*

**DOI:** 10.1101/401604

**Authors:** Michael A. Rieger, Dana M. King, Barak A. Cohen, Joseph D. Dougherty

## Abstract

CELF6 is a RNA-binding protein in a family of proteins with roles in human health and disease, however little is known about the mRNA targets or *in vivo* function of this protein. We utilized CLIP-Seq to identify, for the first time, *in vivo* targets of CELF6 and identify hundreds of transcripts bound by CELF6 in the brain. We found these are disproportionately mRNAs coding for synaptic proteins. We then conducted functional validation of these targets, testing greater than 400 CELF6 bound sequence elements for their activity, applying a massively parallel reporter assay framework to evaluation of the CLIP data. We also mutated potential binding motifs within these elements and tested their impact. This comprehensive analysis led us to ascribe a previously unknown function to CELF6: we found bound elements were generally repressive of translation, that CELF6 further enhances this repression via decreasing RNA abundance, and this process was dependent on UGU-rich sequence motifs. This greatly extends the known role for CELF6, which had previously been defined only as a splicing factor. We further extend these findings by demonstrating the same function for CELF3, CELF4, and CELF5. Finally, we demonstrate that the CELF6 targets are derepressed in CELF6 mutant mice *in vivo*, confirming this new role in the brain. Thus, our study demonstrates that CELF6 and other sub-family members are repressive CNS RNA-binding proteins, and CELF6 downregulates specific mRNAs *in vivo*.

## Introduction

Messenger RNAs (mRNA) are regulated by RNA-binding proteins (RBPs) in every aspect of their life cycle from early steps including transcription, splicing, nuclear export, to later steps such as localization, maintenance, translation into protein, and finally degradation (1, 2). The CUGBP and ELAV-like Factor (CELF) RBP family contains 6 proteins (CELF1-6) which can be divided into two subgroups based on sequence homology: CELF1 and CELF2 which are expressed ubiquitously, and CELF3-6 which show enriched expression in the CNS (3). The most deeply studied, CELF1, was originally characterized in relation to the pathogenesis of myotonic muscular dystrophy (4). It binds CUG-repeat containing sequences and (U)GU-rich motifs (5, 6), and is a multifunctional RBP: CELF1 has been shown to promote both exon skipping (6), and mRNA degradation via recruitment of deadenylation machinery (7). CELFs 3-6 however, have not been as well characterized.

Our laboratory identified CELF6 as both enriched in serotonin producing neurons and disrupted in an individual with autism (8). Upon further study, CELF6 also showed enriched expression in the hypothalamus and in several monoaminergic cell populations commonly targeted for treatment in psychiatry (9). Functionally, CELF6 has been shown capable of regulating splicing *in vitro* (10), however its targets and function *in vivo* are uncharacterized. Here, we were interested in understanding the role of the RBP CELF6, and focused on the brain as the tissue where we had the best understanding of both its expression and evidence that the protein was functional, as germline deletion in mice led to reductions to communicative behavior and other behavioral deficits likely mediated by the brain (8).

In order to define the function of CELF6 *in vivo*, we performed cross-linking immunoprecipitation and sequencing (CLIP-Seq) and found CELF6 to be primarily associated with 3’UTRs of target mRNAs, many of which are involved in synaptic transmission. These sequences showed enrichment for UGU-containing motifs, consistent with previous research on other CELFs, and validating *in vivo* CELF6 binding preferences from cell free systems (11). To comprehensively define the function of CELF6’s interaction with these sequences, we cloned over 400 UTR elements found under CLIP-Seq peaks into the 3’UTR of a reporter construct and measured reporter library expression and ribosomal abundance with and without CELF6 overexpression, and with or without mutation of motif sequences. We found that CELF6 functioned as a repressor of ribosome occupancy and protein production by destabilizing mRNAs containing the wild-type motif sequences, and that this was abolished by motif mutation. We also found that this function was similar across CELF3-6. Finally, we show CELF6 targets are generally derepressed in brains of CELF6 knockout mice, indicating this role is conserved *in vivo*. Taken together, we show that CELF6 and other CELF family members can repress transcript abundance of key mRNAs, and thus may fulfill an important role in regulating cellular function in the brain.

## Results

### Celf6 primarily associates with 3’UTRs of target mRNAs *in vivo*.

To define the *in vivo* binding locations of CELF6, we performed CLIP on brains from BAC transgenic mice expressing an epitope tagged CELF6-YFP/HA (78 kDa) with the endogenous CELF6 pattern (9). As CELF6 molecular function has not been studied *in vivo*, we first confirmed CELF6 binds RNA in the brain. We performed CLIP with anti-EGFP antibodies on CELF6-YFP/HA mice followed by radiolabeling of nucleic acid (Figure 1A) across several RNase concentrations. Controls included an immunoprecipitated (IP) from uncrosslinked samples and IP from wild-type (WT) tissue. As expected, there was a lack of detectable RNA in IP from WT tissue and uncrosslinked YFP+ tissue. Next, to capture the targets of CELF6 *in vivo*, we chose a region approximately 60-200 nucleotides in size (80-150 kDa) to purify from lysates of 4 pools of CELF6-YFP/HA+ brains. Similar to Vidaki et al.’s study of Mena (12), we found that the stringent lysis and wash conditions of standard CLIP protocols were incompatible with CELF6 IP (not shown). Therefore, to enable rigorous statistical definitions of CLIP targets, we also collected 4 replicates of CLIP samples generated from pools of 3-4 brains from 4 independent litters, and a key additional control:. IP from 3 pools of WT littermate brains to identify RNAs that interact non-specifically with the capture reagents. Likewise, some CLIP studies fail to account for differences in RNA abundance in the starting input RNA, thus biasing their results towards highly expressed genes (13). We therefore also collected 2% input samples to measure starting transcript abundance and to allow statistical identification of mRNAs bound by CELF6.

**Figure 1.**
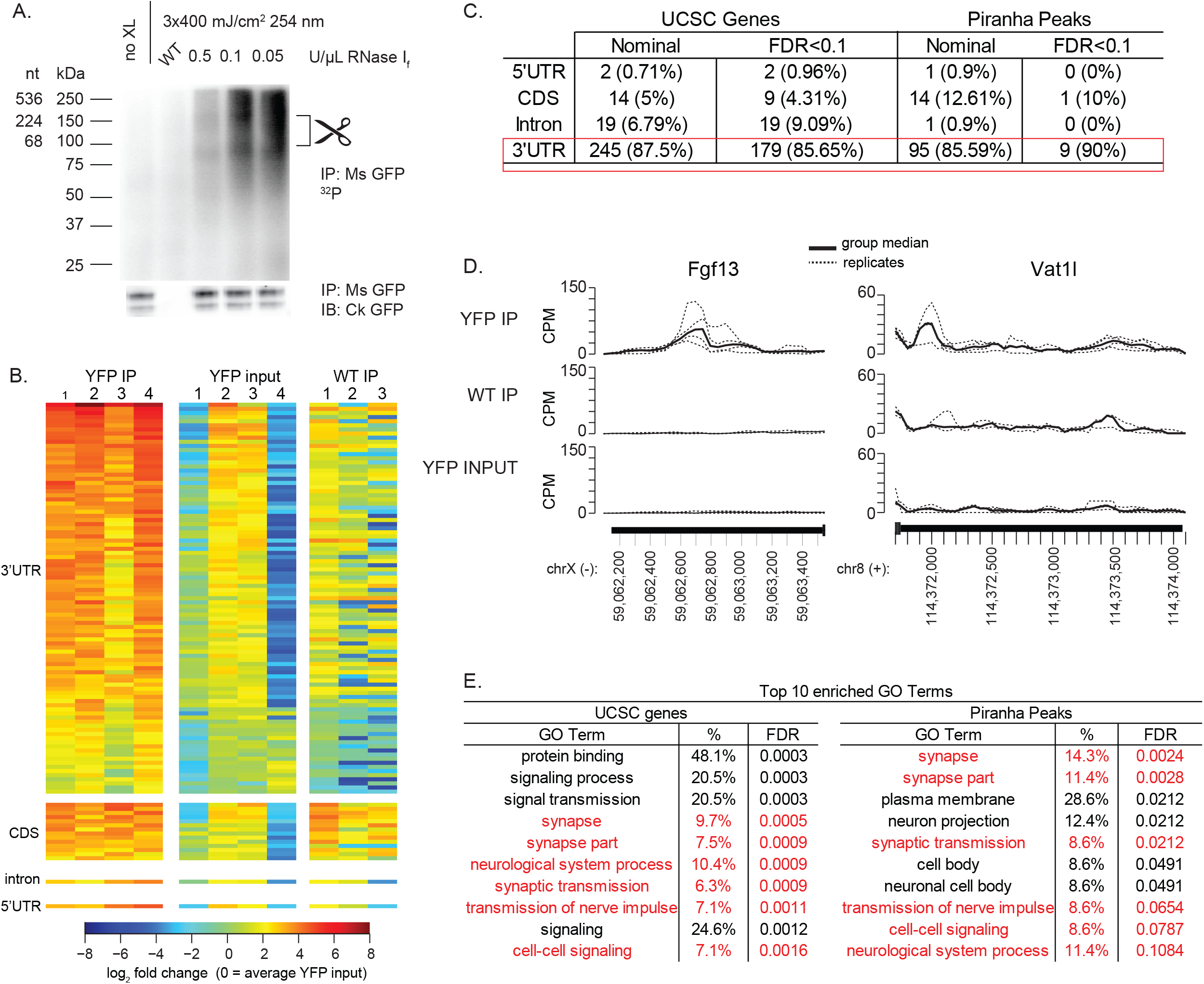
Celf6 primarily associates with 3’UTRs of target mRNAs *in vivo*. (A) CLIP and 32P labeling of bound RNA. Each lane is one brain worth of material for no crosslink(no XL), wildtype(WT) control, and 3 RNase I concentrations for digestion. Upper panel: autoradiogram of end labeled RNA. Bottom panel: immunoblot of anti-GFP showing CELF6-HA/YFP. The scissors mark region (80-150 kDa) above CELF6-HA/YFP (78 kDa) isolated for sequencing. Immunoblot detects two bands, corresponding to sizes of both known isoforms of CELF6 when tagged with HA/YFP. (B) log_2_ counts-per-million (CPM) RNA in 4 CELF6-HA/YFP+ replicates and 3 WT replicate samples across nominally significant regions identified by Piranha peak calling followed by edgeR differential enrichment analysis. Heatmaps show enrichment in HA/YFP+ immunoprecipitate (IP) samples relative to input and WT controls. CPM normalized to mean YFP input by row. (C) Summary of differential enrichment analysis of genes showing nominally significant (p<0.05), and Benjamin-Hochberg (FDR<.1) enrichment in HA/YFP+ IP samples relative to both HA/YFP+ input and WT IP controls, for both the UCSC Gene Annotation method as well as Piranha peak calling method. No peaks were found in intergenic regions. (D) Example traces for two identified CLIP target 3’UTRs, showing CPM in YFP IP samples compared to controls. (E) BiNGO analysis for gene ontology (GO) terms enriched in CLIP target genes determined according to either differential analysis methods.

We prepared sequencing libraries from these samples using an adaptation of the eCLIP workflow (14) (Supplemental Protocol S1). All samples were sequenced to a similar depth, and on average 94% of all uniquely aligning reads mapped to genic regions, consistent with RNA expression patterns (Supplemental Table S1). Next, to define specific sites of CELF6 binding, we called peaks throughout the genome using Piranha (15) and summed reads under these peaks. We then performed differential enrichment analysis comparing CLIP to controls using edgeR (16) (Supplemental Table S2). In case binding was more general across a transcript, we also summed reads mapping to subgenic regions: 5’ untranslated (5’UTR), coding sequence (CDS), introns, or 3’ untranslated (3’UTR) of annotated genes and performed differential enrichment analysis (Supplemental Table S3). In total, we identified significant enrichment across 364 genes combined across analysis methods.

CELF proteins have been identified to function in alternative splicing of mRNAs as well as post-transcriptional regulation. To gain insight into the molecular functions of CELF6 *in vivo*, we examined where CLIP reads were enriched relative to subgenic region (5’UTR, 3’UTR, introns, CDS). We hypothesized that CELF6 splicing-related functions would correspond to increased density in internal coding exons and alternatively spliced introns, while post-transcriptional regulatory functions would show increased density in UTRs. Figure 1B shows a heatmap of abundance in all samples under significant Piranha peaks relative to controls, and indicates that the vast majority of differentially enriched peaks are in 3’UTR regions. This was true in both in both analytical approaches (Figure 1C, >85% of all CELF6 bound regions). Figure 1D shows example read distribution in CPM across the 3’UTRs for two high confidence targets, *Fgf13* and *Vat1l*. The *Fgf13* gene is part of the FGF-like family of genes, and controls localization of voltage-gated Na+ channels in axons and *Vat1l* (Vesicle Amine Transport 1-Like) is a paralog of the *Vat1* gene which regulates monoaminergic neurotransmitter storage and release (Eiden et al., 2004; Pablo et al., 2016). Overall, a Gene Ontologies analysis revealed that CELF6 targets were disproportionately found to be involved in synaptic transmission (Figure 1E). Together with binding in the UTR, CELF6 regulation of such genes may thus directly impact neuronal cell function by regulating the stability or translation of targets.

### CELF6-bound 3’ UTRs are enriched for U-rich and UGU-, CU- containing motifs

Enriched binding in 3’ UTRs suggests CELF6 has affinity for specific nucleotide motif sequences *in vivo*, as previously reported in biochemical assays on recombinant protein (RNACompete, (11)). Thus we next defined the sequence specificity of CELF6 binding. Manual inspection of sequences under peaks found many matched the UGU-rich preferences identified via RNACompete (Figure 2A). However, to systematically identify motifs enriched in CELF6 3’UTR targets we used the MEME suite tools to identify both *de novo* motifsand previously cataloged motifs via Analysis of Motif Enrichment, AME based on the CISBP-RNA database (Supplemental Table S9: Key Resources). We analyzed 50 nucleotide segments centered under 491 peak maxima. As some UTRs possessed more than a single peak, these represented 174 unique UTRs. As a control, we chose 491 sequences sampled randomly from the 3’UTRs of brain expressed genes showing no evidence for CELF6-YFP/HA binding.

**Figure 2.**
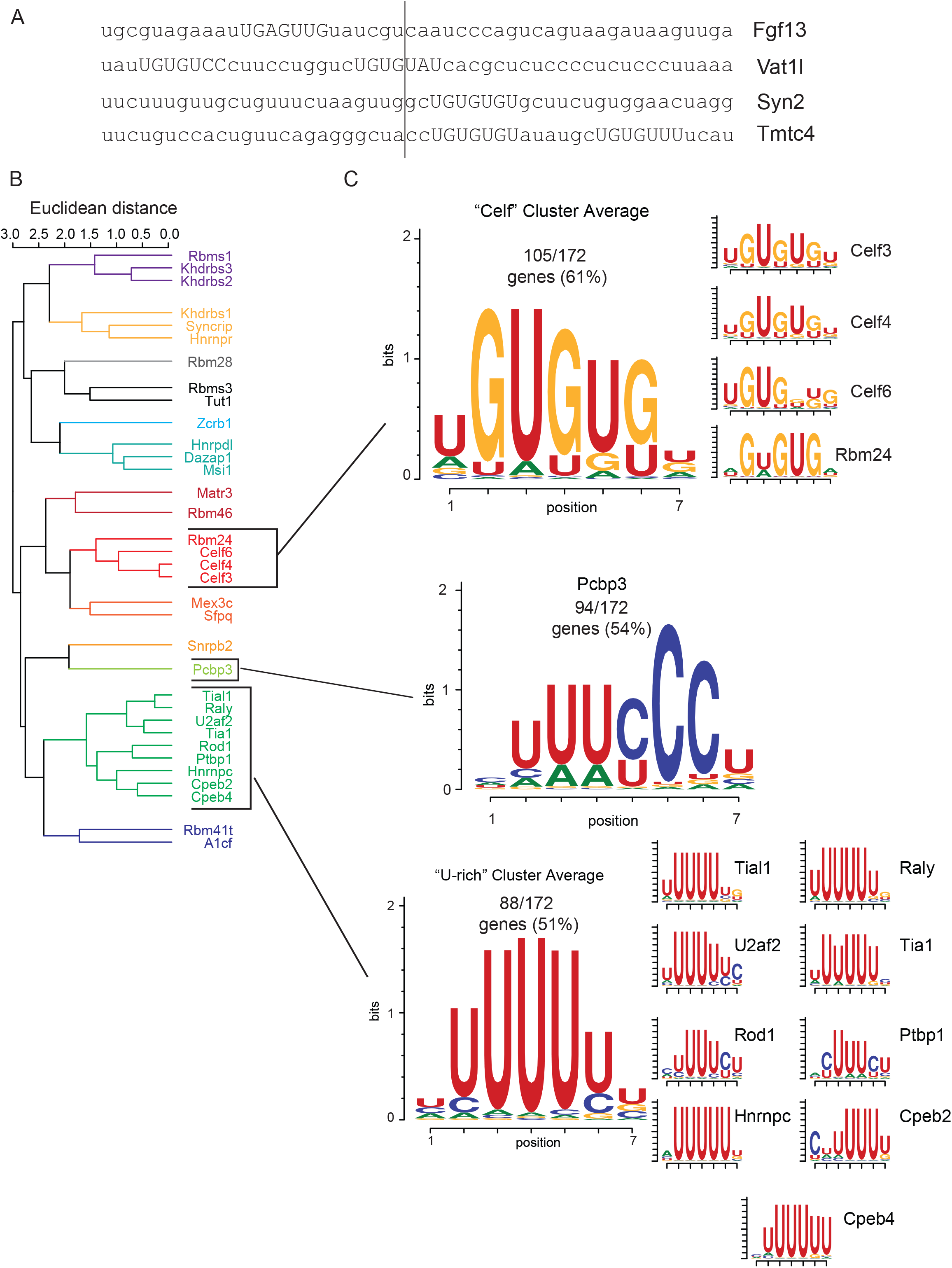
CELF6-associated 3’UTRs are enriched for U-rich and UG-, CU- containing motifs. (A) Example 50-nt regions under CLIP peaks in the 3’UTRs of Fgf13, Vat1l, Syn2, and Tmtc for, all showing evidence for UG-rich sequence. (B) Clustering of Analysis of Motif Enrichment (AME) identified motifs in CELF6 CLIP peak regions compared to randomly sampled control genes measurable in background samples (HA-YFP input). Color coding shows cluster membership. (C) Motif logos for the common clusters found in CLIP target peak regions. Logos represent the average of position weight matrices (PWMs) for each motif cluster, as well as the individual PWMs making up each cluster’s membership.

We found 36 enriched motifs from the CISBP-RNA database at FDR<0.1 (p<0.001, AME Ranksum test on maximum odds score). These included RNACompete binding motifs for CELF3, CELF4, and CELF6, as well as the U-rich motifs of the TIA and CPEB families of RNA binding proteins. RBPs can exhibit degenerate preferences (17, 18). Therefore, to generate a more holistic understanding of CELF6/sequence interaction, we clustered enriched motifs by the Euclidean distance between their position weight matrices (PWMs). As expected, motifs clustered partially by RNA binding protein family (Figure 2B). Motifs within a cluster are highly similar, such as the CELF motifs which all show RNAcompete preferences for UGU-containing sequences. We then scanned our sequences to determine which showed significant matches to the cluster average PWMs. 473 of 491 sequences showed significant matches to at least one cluster.

There were a small number of motifs that were found recurrently across the sequences (Figure 2C, Supplemental Table S4). 105/172 unique genes’ 3UTRs (61%) showed at least one match to the “CELF”- cluster ([U/A]GUGU[G/U][UGA]). In addition to the CELF cluster, 94/172 genes’ 3’ UTRs (54%) showed at least one match to the PCBP3 motif which forms its own cluster possesses a central UUU[C/U]CC sequence. PCBP3 can bind both double stranded and single stranded nucleic acid and is known primarily as a transcription factor (19) though a related protein (PCBP4) has also been shown to regulate mRNA stability (20). 88/172 (51%) genes showed at least 1 match to the “U-rich” cluster, whose members all possess a central stretch of 4-5 Us. Proteins with binding motifs in this cluster include TIA1, which is involved in stress granule localization (21), CPEB proteins, which are known to be involved in polyadenylation (22), and HNRNPC, which is involved in both mRNA stability and localization/nuclear trafficking (23, 24). 59/172 genes (34%) had at least one match to both the CELF cluster & the U-rich cluster, and 72/172 genes (42%) showed at least one match to both the CELF cluster and PCBP3. Thus, CELF6 binding appears to be mediated by specific sequences that are combinations of known motifs for both CELFs and other RBPs.

### Massively Parallel Reporter Assay defines the function of CELF6 bound motifs

Our motif analysis indicates sequence specificity mediates the interactions between CELF6 and RNA *in vivo*, but the downstream consequence of CELF6 association to elements in the 3’UTR of mRNAs has not been defined. Because 3’ UTR elements often regulate protein production, we first sought to assess the function of these sequences or ribosome occupancy. Rather than focus on a single or small number of selected elements for analysis, which might or might not be representative, we utilized a massively parallel reporter assay for post-transcriptional regulatory elements (PTRE-Seq) to evaluate all of them (25). For PTRE-Seq, we sub-cloned 436 independent CLIP-defined UTR elements, each 120 bp long and centered under CLIP peaks, into the 3’ UTR of a tdtTomato expression plasmid (Figure 3A). To cast a wide net for potential targets, this comprised any clonable element with a nominally significant enrichment by CLIP (Supplemental Table S5). To ensure reproducibility, each UTR element was included in the library design 6 times, with a unique 9-bp barcode to provide internal replication and buffer against barcode effects. In addition, to assess candidate motifs within each element, we also included 436 matched elements with all significant motif matches mutated for comparison with the unmutated (reference) sequences (see Materials and Methods: PTRE-Seq Reporter Library Preparation).

**Figure 3.**
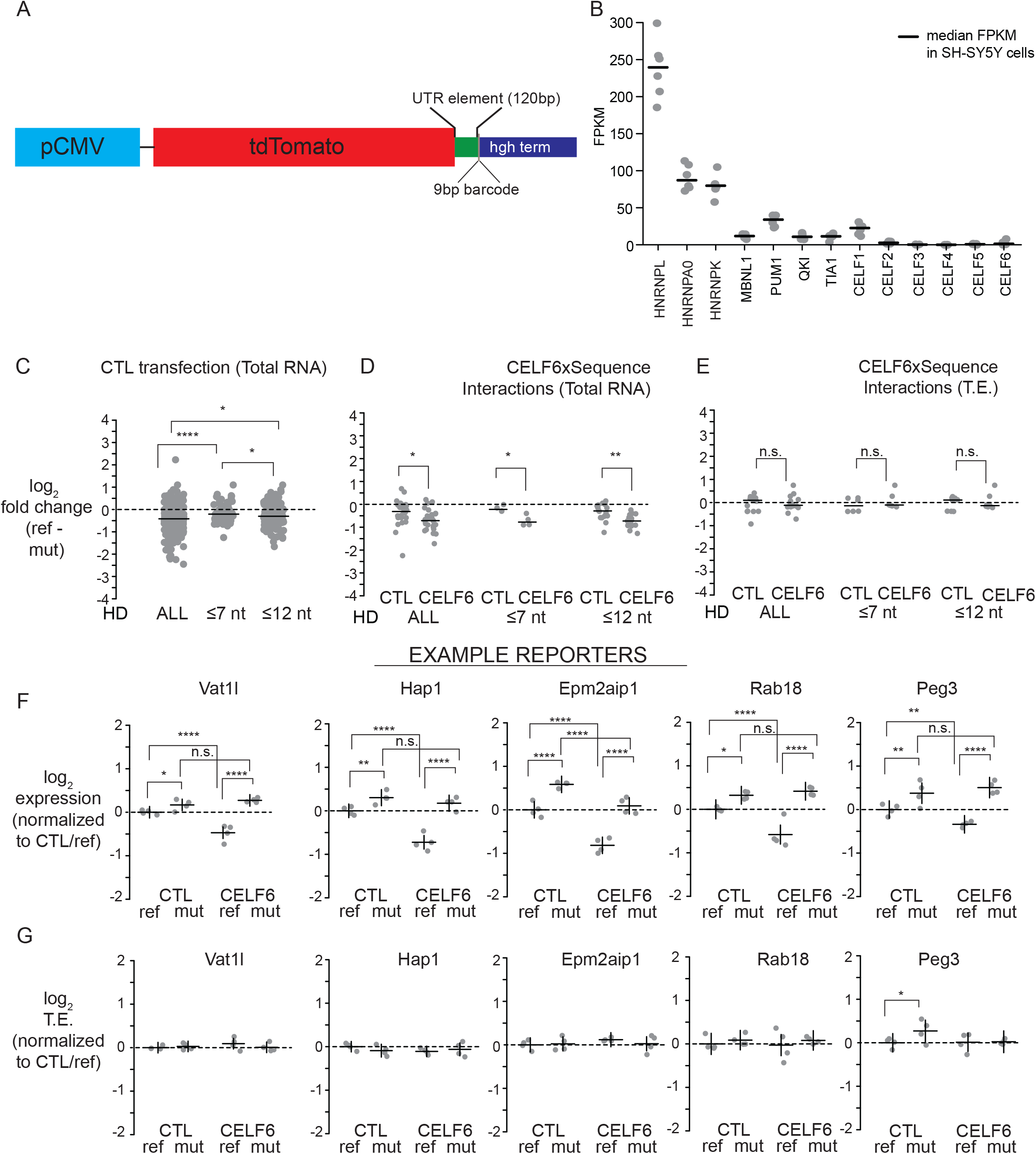
CELF6 CLIP enriched motif sequences represent a set of repressive elements. (A) PTRE-Seq tdTomato reporter (pmrPTRE_AAV). Expression of mtdTomato (membrane-localizing tdTomato)is driven by a CMV promoter. 120-nt regions under CLIP peaks, or the same sequence with motifs mutated, are subcloned after the tdTomato stop codon followed by a 9-nt barcode sequence, and a human growth hormone polyadenylation signal. (B) Fragments per Kilobase of transcript per Million reads (FPKM) levels are shown for several RNA binding proteins in RNA extract from 8 replicate platings of SH-SY5Y cells. (C) Recovered reporter RNA log_2_ fold changes in expression between reference and mutant elements in the CTL transfection conditions, considering all reference-mutant pairs, as well as only reference-mutant pairs possessing ≤7 nucleotide subsitutions, or ≤12 nucleotide subsitutions. (D) Recovered reporter RNA log_2_ fold changes in expression between reference and mutant pairs in CTL and CELF6 overexpression conditions from elements showing significant condition X sequence interactions. (E) Recovered reporter RNA log_2_ fold changes in translation efficiency (TE) between reference and mutant pairs in CTL and CELF6 conditions for elements showing significant condition X sequence interactions. Data points in (C)-(E) are average estimates of log_2_ fold change reference vs. mutant per condition, averaged by replicate over barcodes, and then averaged over replicates, with lines representing medians for each distribution. Comparisons between conditions or sequence mutation were assessed by Mann Whitney U tests. (F) Log_2_ expression across 5 example reporter library elements in CTL, CELF6, reference and mutant conditions. (G) Log_2_ TE across example reporters in (F). Data points in (F)-(G) are averaged across barcodes, with each a biological replicate. Horizontal lines represent average (normalized to the CTL/reference sequence condition). Vertical lines represent 95% confidence intervals. Post hoc pairwise comparisons between conditions shown were computed using the multcomp package in R with simultaneous multiple comparisons corrections using the multivariate normal distribution (single step method in multcomp). Significance notation: n.s. p>0.1, † p<0.1, *p<0.05, ** p < 0.001, *** p<0.0001, **** p<1E-5.

Next, we assayed the impact of each element on final translation levels, as assessed by ribosome occupancy of these reporters. We used transient cotransfection of the library with an EGFP/RPL10A construct that tags ribosomes with GFP enabling Translating Ribosome Affinity Purification (TRAP). This method allows assessment of RNA abundance on ribosomes by GFP pull down, and has previously been shown to be sensitive to UTR elements that regulate translation (Heiman et al., 2008) (Supplemental Protocol S2). As a control we collected total RNA from the same cells. We used SH-SY5Y neuroblastoma cells as a model system as total RNA-Seq data revealed that, with the exception of *CELF1*, *CELF*s 2-6 were low or largely undetectable (Figure 3B). Thus, we could also use this system to control *CELF6* levels by adding human *CELF6* exogenously via co-transfection with an His/Xpress epitope tagged *CELF6* construct (Ladd et al., 2004) or empty pcDNA3.1 vector (CTL), with overexpression of *CELF6* RNA confirmed by RTPCR (Supplemental Figure 1). Finally, to determine the impact of each element on ribosome occupancy in response to these manipulations, we prepared RNAseq libraries from all TRAP and total RNA samples (Supplemental Protocol S3), and quantified each barcode’s abundance. All conditions were replicated on four independent days to ensure robustness and reproducibility. After normalization (below) this design enables us to identify elements that potentially alter protein translation, and assess their response to CELF6 expression with statistical confidence.

Prior to statistical testing for experimental effects, we first verified the reproducibility of replicates prepared with this approach. The Pearson correlation in elements counts between TRAP replicates was between 0.93 and 0.95, and 0.92 and 0.97 for total RNA controls, indicating good reproducibility. Next, to account for any differences in starting abundance of library members, log_2_ CPM in RNA barcodes were normalized to log_2_ CPM from sequencing the starting DNA plasmid pool (“log_2_ expression”). After removing barcodes which were absent or poorly detected across all samples, the final analyzed library contained 423 UTR element pairs (reference and mutant) all of which were represented by 3-6 barcodes per element, across 172 total genes.

We then applied statistical models to identify effects of CELF6 presence and its interaction with motif sequence to alter ribosome occupancy as assessed by TRAP. Specifically, to determine the relationship between CELF6 overexpression and element sequence, we analyzed log_2_ TRAP levels using a 2x2 factorial design linear mixed effects model, fitting fixed effects of element sequence (“reference” or “mutant”) and overexpression condition (CTL or CELF6 overexpression) and the interaction of condition X sequence, with a random intercept term for each element, treating individual barcodes as a repeated measure. Individual models were fitted for each UTR element pair. A summary of effects across all library elements and estimates of R^2^ is shown in Supplemental Table 6. At nominal p<0.05, 332/423 (78.5%) of elements showed a significant effect (main effects of either sequence or condition, or sequence X condition interaction, 327/423(77.3%) at FDR <0.1).

### CELF6 bound motif sequences repress ribosome occupancy by decreasing RNA abundance.

We first examined the role of the element sequence itself on ribosome occupancy by comparing the UTR elements with and without motif mutations in the TRAP data. Among the 78.5% of elements which showed significant effect (sequence, condition, or interaction), 88.6% (294 elements) showed a main effect of sequence regardless of CELF6 expression. Looking at the distribution of log_2_ fold changes in expression between reference and mutant sequence, 294 elements. 247/294 (84%) of these were repressive when compared to their mutated counterparts (median log_2_ fold change -0.48, 1.39-fold decrease). Even looking at estimates of fold change in CTL conditions across all 423 elements, regardless of significant effects, 305/423 (72.1%) had negative values. Thus CELF6 bound elements were repressors of ribosome occupancy, even in the absence of CELF6 overexpression

There are two primary mechanisms by which a sequence can alter ribosome occupancy – altering translation efficiency (TE), usually reflecting the loading of mRNAs onto ribosomes, or altering mRNA stability (26–29). To assess TE, log_2_ expression for each barcode in TRAP samples was normalized to input RNA to account for their overall abundance (log_2_ TE). We then we fitted our model independently for both input RNA expression, to assess abundance, and TE. A summary of effects across all library elements and estimates of R^2^ is shown in Supplemental Tables 7 & 8 (input expression and TE, respectively) along with Benjamini-Hochberg FDR across all elements. At nominal p<0.05, 336/423 (79.4%) of elements showed any significant effect on either TE or input expression (312/423, 73.8% at FDR <0.1), indicating the majority of sequences were sensitive to mutation, CELF6 overexpression, or both.

To assess the relative influence of either mechanism on ribosomal occupancy effects, we first looked at input RNA expression. 316/423 elements (74.7%) showed any significant effect (312/423, 73.8% at FDR<0.1). Similar to what was found in analysis of TRAP RNA levels, 91.1% of these (288/316) showed main effects of sequence mutation, regardless of CELF6 overexpression. As with ribosome occupancy, 84.7% had fold changes less than 0 indicating that reference sequences are repressive when compared to their mutated counterparts (Figure 3C) (median log_2_ fold change -0.43, 1.35-fold decrease). However, unlike input RNA expression, we did not observe this trend in repression in TE. Only 89/423 sequences showed any nominally significant effect on TE and 0 showed any significant effect at FDR <0.1, 48 of which (54%) showed main effects of sequence. 18/48 (37.5%) showed log_2_ fold changes in TE less than 0, with a median value of 0.12 (1.09 fold change) This indicates that the mechanism for the change in ribosome occupancy is primarily alteration of transcript abundance, rather than TE.

As UTR elements with larger numbers of motif matches have higher numbers of mutated bases, one concern is that mutating these motifs may have altered a large fraction of the element rather than just a core motif. Thus to mitigate this concern, we subsetted our data to look at changes between reference and mutant for elements with smaller numbers of mutated bases (Hamming distance (HD) of ≤7 or ≤12 nucleotides). Expression of the reference sequences were still generally lower than mutant sequences for these subsets, with 71.6% less than zero and 77.8% less than zero for HD ≤7 and HD ≤12 respectively. The median log_2_ fold change for HD≤7 was -0.21 (p=4.2E-6, Mann-Whitney U test compared to all elements, N=67 HD≤7) and the median log_2_ fold change for HD≤12 was -0.30 (p=0.0011 Mann-Whitney U test compared to all elements, N= 189 HD≤12, p=0.022 compared to HD≤7). Thus even a small number of mutations is capable of elevating the expression levels of most elements.

### CELF6 enhances repression by decreasing RNA abundance in a sequence dependent manner

We next assessed interactions between CELF6 overexpression and sequence, focusing on RNA expression. 30 elements (29 unique genes) showed a nominally significant interaction (p<0.05) between CELF6 overexpression and sequence. Figure 3D plots the average log_2_ fold change between reference and mutant sequence for both the CTL and CELF6 conditions across each subsetted level of sequence mutation. In each case, the fold change between reference and mutant was more steeply negative in the CELF6 condition (CELF6 median log_2_ fold change reference – mutant -0.71 (~1.6-fold decrease), CTL -0.31, p = 0.001 | HD ≤ 7: CELF6 -0.78, CTL -0.22, p = 0.028 l HD ≤ 12 CELF6 -0.78, CTL -0.29 p = 4.8E-4, Mann Whitney U-tests). This is driven by repression of the reference sequence rather than an elevation of the mutant sequence by CELF6 (Median log_2_ fold change CELF6-CTL reference sequences = -0.38, CELF6-CTL mutant sequences -0.04, p = 2.1E-4 Mann Whitney U-test). This indicates the CELF6 impacts these elements by further decreasing mRNA levels, and this effect is sequence dependent.

As mentioned earlier, few effects were observed on TE by comparison. Out of the 89 elements showing nominally significant effects on TE, only 18 showed interactions with CELF6 overexpression at p<0.05, with 0 showing interactions at FDR < 0.1. In Figure 3E, we plot the log_2_ fold change in TE for these elements. To understand the source of significant interaction in the cases where it was detected, we looked at post-hoc multiple comparisons and overall we did not detect a consistent direction of CELF6 effect on TE across elements. In particular, looking at changes to reference sequence TE when CELF6 is overexpressed, only 4 elements showed a difference compared to CTL. Two of these showed an increase (Fnbp1l p=0.0097, log_2_ fold change 0.43 l Reep1 p=0.004 log_2_ fold change = 0.90), and two showed a decrease (Peg10 p = 0.008 log_2_ fold change -0.37 l Lin7c p=0.006 log_2_ fold change = -0.54). By contrast, looking at input RNA expression, 19 out of 30 elements with nominally significant interactions showed log_2_ fold changes in reference between CELF6 and CTL which were negative. These ranged between -1.45 to -0.22 with a median of -0.55 (≈1.5-fold reduction). Example element expression is shown in Figure 3F and TE in Figure 3G. These findings indicate that for elements showing interactions between CELF6 and element sequence, there is a repression of the reference sequence via a decrease in RNA abundance with CELF6 expression, and that this effect is abolished after mutation of the conserved motifs. Therefore we conclude that, overall, CELF6 decreases RNA abundance. Further, where impact on TE occurs, it is not generalizable in direction of effect.

In order to further validate these findings and confirm an impact on final protein levels, we overexpressed five individual reporters, with and without mutation, along with EGFP-tagged CELF6 or EGFP alone, then measured reporter expression by tdTomato fluorescence (Figure 4). For 4/5 cases, repression upon CELF6 overexpression was the same direction and magnitude to that observed by PTRE-Seq. Furthermore for 3/4, the repression observed by CELF6 was significantly reversed by motif mutation. This confirms the finding in independent assays in shows CELF6 ultimately impacts protein levels of its targets.

**Figure 4.**
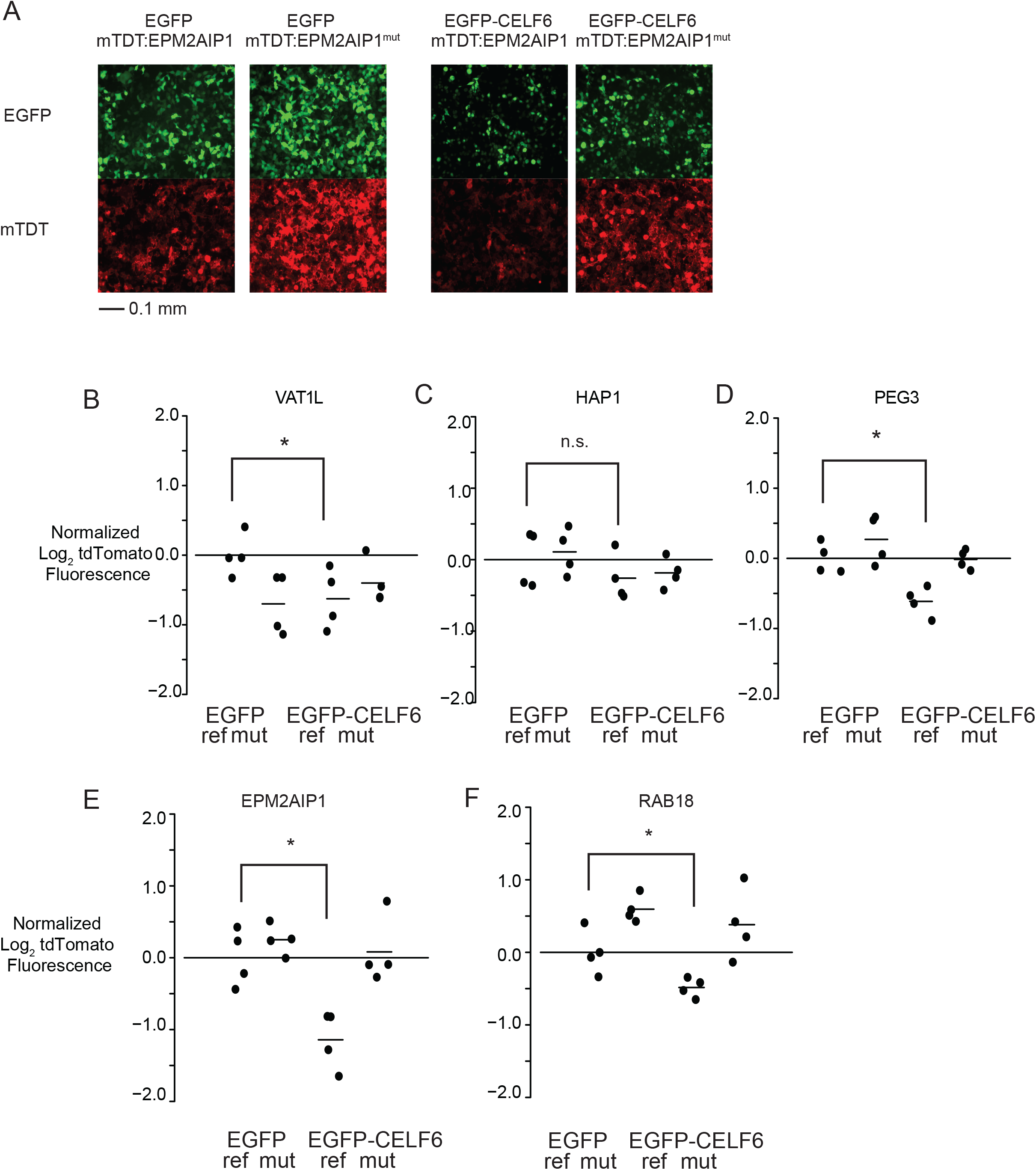
Individual mtdTomato UTR element constructs replicate interactions of CELF6 overexpression and sequence. Individual UTR elements from VAT1L, HAP1, PEG3, EPM2AIP1, and RAB18 were cloned as in Figure 3A, and overexpressed along with either EGFP or EGFP-CELF6. (A) Example live cell epifluorescent images from a transfection experiment showing EGFP or EGFP-CELF6 expression (upper panels) and either tdTomato:EPM2AIP1 or tdTomato:EPM2AIP1(mutant) (lower panels). (B)-(F) tdTomato log_2_ fluorescence, normalized to EGFP/reference sequence UTR element sample mean. Two-way ANOVA was computed for each element with significance shown for pairwise post-hoc comparisons with simultaneous correction using the multcomp package in R. n.s. p>0.1, † p<0.1, *p<0.05, ** p < 0.001.

### CELF3-5 also enhance repression via decreasing RNA abundance.

CELF3, CELF4, and CELF6 binding preferences determined by RNAcompete are highly similar (Figure 2), and as a group, CELF3-6 are more similar in amino acid identity than CELF1 or CELF2 (3). Therefore, we hypothesized that the repression of elements we observed for CELF6 would also be true of CELF3-5. Thus we transiently transfected our PTRE-Seq library along with His/Xpress-tagged human CELF3, CELF4, or CELF5 used previously to study these proteins (Supplemental Figure S1) (10, 30). We then performed the analysis of mRNA abundance described in the preceding section, refitting the linear mixed models to include data from these new overexpression conditions (Figure 5).

**Figure 5.**
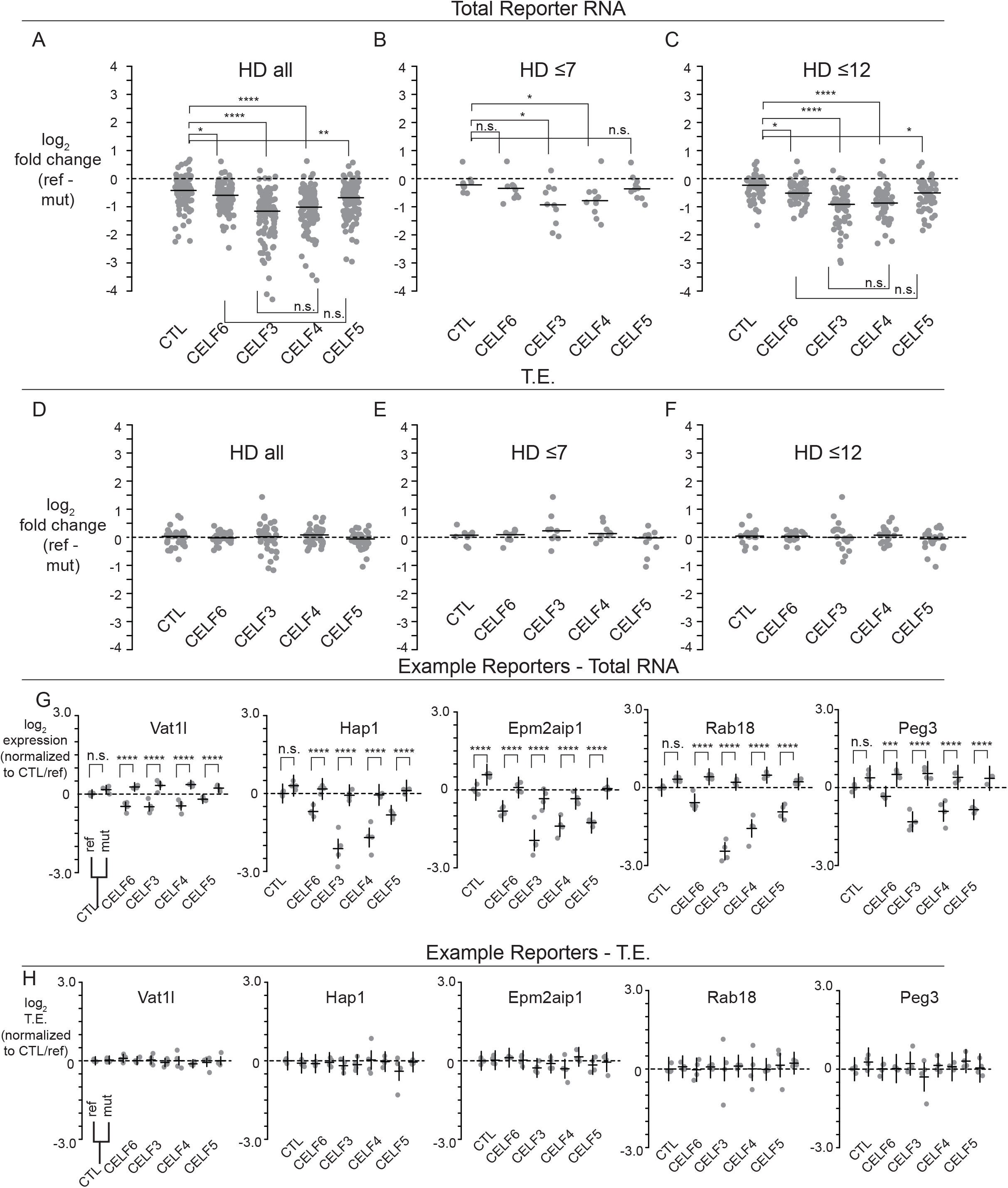
CELF3-5 show redundancy in ability to enhance repression of CELF6-CLIP enriched UTR elements. (A)-(C)Log_2_ fold change across CELF conditions considering all reporter elements, or reporter elements with a difference of ≤7 or ≤12 nucleotides between reference and mutant sequences, for any element with significant condition X sequence interactions. (D)-(F) Log_2_ fold change in TE across conditions as in (A)-(C). Data points in (A)-(F) were averaged for redundant barcodes, then across replicates, with lines representing medians. Statistical comparisons in (A)-(F) were assessed by Mann Whitney U tests. (G) Log_2_ expression across 5 example reporter library elements across CELF, reference and mutant conditions. (H) Log_2_ TE across example reporters in (G). Data points in (G)-(H) are averaged across redundant barcodes, with each dot representing a replicate. As in Figure 3, horizontal lines represent average expression or TE (normalized to the CTL/reference sequence condition), and vertical lines represent 95% confidence intervals. Multiple comparisons corrections computed with multcomp in R. n.s. p>0.1, † p<0.1, *p<0.05, ** p < 0.001, *** p<0.0001, **** p<1E-5.

There were 111/423 (p<0.05, 91/423 FDR < 0.1) elements representing 89/172 genes showing nominally significant sequence by CELF expression interactions in total reporter RNA. Looking at the log_2_ fold change in reference vs. mutant sequence, CELF3 and CELF4 showed the largest differences (Figure 5A). The median element responding to CELF3 expression shows a -1.15 log_2_ fold lower expression of the reference sequence compared to the mutant sequence. CELF4 was comparable with a median -1.01 log_2_ fold change (CTL -0.42, CTL vs. CELF3 p = 3.5E-15, CTL vs. CELF4 p=1.5E-12, CELF3 vs. CELF4 p = 0.13, Mann Whitney U tests). CELF6 and CELF5 also showed comparable fold changes, and these were intermediate between the CTL condition and CELF3/CELF4 (CELF5: -0.68, CELF6: -0.59, CELF6 vs. CTL p=0.0024, CELF5 vs. CTL p=0.00034, CELF5 vs. CELF6 p = 0.26) (Figure 5B). Thus overexpression of any of these CELFs was able to reduce abundance of reference reporters, an effect which could be abolished by mutation. Furthermore, CELF3 and CELF4 were associated with the strongest effects, and CELF5 and CELF6 showed more moderate effects. Thus within these CELF proteins, CELFs 5/6 and CELFs 3 /4 appear to form distinct subgroups with respect to their effects on mRNA abundance.

When looking at TE, there were fewer significant effects. 41 elements showed significant interactions of condition and element sequence (p<0.05, 6/423 at FDR<0.1). However, when plotted in terms of their fold change between reference and mutant sequence, we were again unable to generalize a direction of effect on TE across elements. Median log_2_ fold changes in TE were near 0 with no significant differences across conditions (Figure 5D-F) Thus CELF overexpression decreased overall ribosome occupancy of reference elements, but did so largely by disrupting reporter transcript abundance. Total reporter expression for key examples are shown in Figure 5G with their respective TE shown in Figure 5H.

As expression of various CELFs may overlap in the brain, we next tested if co-expressing CELF proteins exerted additive or synergistic effects on reporter expression. We transfected CELF6 construct with one of CELF3, CELF4, or CELF5. Among elements showing significant sequence by condition interactions, overexpression of CELF3 and CELF6 together resulted in repression similar to CELF3 by itself (Median CELF3/CELF6 log_2_ fold change compared to CTL: -0.95, CELF3 alone -0.86, p = 0.38). This was also true of CELF4 (Median CELF4/CELF6 log_2_ fold change compared to CTL: -0.85, CELF4 alone -0.62, p=0.14). Thus the effect of CELF4 and CELF3 appears to be dominant, or at least maximal, in these co-transfections. We confirmed this is not due to differences in the level of CELF overexpression (Supplemental Figure 1). When CELF5 and CELF6 were expressed together, the median log_2_ fold change compared to CTL was approximately doubled. (Median log_2_ fold change CELF5/CELF6 vs. CTL: -0.48, CELF6 vs. CTL -0.24, CELF5 vs. CTL -0.16, p CELF5/CELF6 vs. CELF5 alone = 0.03), indicating these two may act additively. Thus, overall our reporter assays show that CELF proteins can function as translational repressors of varying magnitudes by suppressing mRNA abundance of targets containing specific sequence motifs.

### CELF6 regulates transcript abundance in the brain

Finally, we sought to determine whether CELF6 also has the same impact on transcript abundance *in vivo* by deleting CELF6 in the mouse brain. We examined the expression of all CLIP-defined targets, using microarray analysis comparing 8 WT and 8 CELF6 knockout (KO) brains. In the microarray data, both probes measuring *Celf6* are clearly reduced, confirming loss of transcript via nonsense mediate decay (Figure 6A). Aside from this, consequences of CELF6 loss on total brain RNA abundance are fairly modest: while several hundred genes show a nominally significant change in RNA abundance (Figure 6B) the median log_2_ fold change is 0.24, with only the Celf6 probes surviving genome-wide multiple testing correction. However, examining the distribution of fold changes across all CLIP targets, compared to a similar set of randomly selected probes, clearly reveals that the CLIP targets are significantly more abundant in KO mice(p< 1.11e-09, Welch’s T-test) (Figure 6C), consistent with a role for Celf6 in decreasing specific messenger RNA stability *in vivo*. Given that only limited cells in the brain express CELF6 (9) and many may also have compensatory expression of other CELF family members, modest overall changes are unsurprising. Likewise, this experiment assesses both direct effects on CELF6 targets and indirect effects on gene expression due to a lifetime of development and function in the absence of CELF6. However, in spite of these two limitations on sensitivity, we see nominally significant (p<0.05) regulation of 21 of the CELF6 CLIP targets including Fos, Mecp2, Reln and Fgf13 (Figure D). All but one of these 21 CELF6 targets change in the expected direction, showing increased mRNA abundance with the loss of CELF6, a result highly unlikely to be due to chance (p<.0005, χ^2^Test). This is driven by the 3’UTR CLIP targets, (p<2.2e-08, Figure 6C), as the identified targets in 5’UTR and introns show no median change. CDS sequences show the same magnitude of increase as 3’ UTR elements with CELF6 KO, though N is less and this is not significant. Thus, CELF6 deletion generally causes an increase in the RNA abundance of its targets, *in vivo*, as it does in reporter assays *in vitro*.

**Figure 6.**
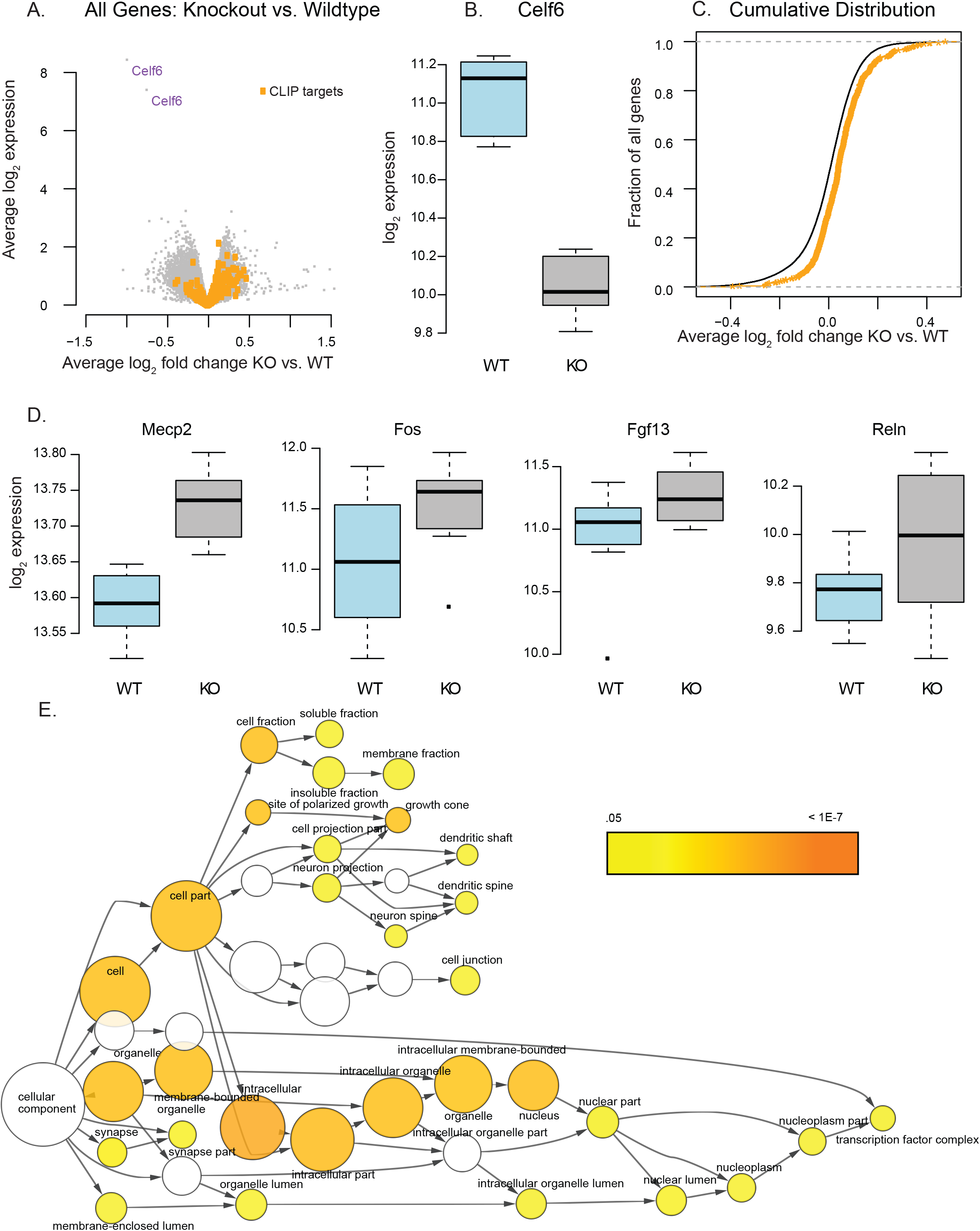
CELF6 targets identified by CLIP show increased expression in Celf6 knockout brains. A) Volcano plot shows differences in gene expression between wildtype and CELF6 knockout(KO) mouse brain. B) Examination of CELF6 reveals KO has decreased CELF6 expression, as expected (p<10E-6). C) CLIP targets (orange) are de-repressed in KO brains when compared to other genes (p<1.1E-9, Welch’s T-Test). D) Examples of CLIP targets with higher mRNA abundance (p<.05) in KO mouse brains include *Mecp2, Fos, Fgf13*, and *Reln*. E) Gene Ontologies analysis of upregulated mRNAs after Celf6 knockout reveals a disproportionate number of neuronal projection localized proteins, as well as nuclear transcription factors.

An exploratory systems analysis of the consequences of CELF6 loss is also intriguing. Gene Ontologies analysis of the 200 most upregulated transcripts (p<.05) regardless of CLIP status reveals significant enrichment for terms for transcription regulators (transcription factor complex, p<.8E-3 with B-H correction) and other components of the nucleus on one hand, and synaptic proteins on the other (growth cone, p<3.47E-4, and dendritic spine (4.02E-3) (Figure 6E). Examination of the down-regulated genes showed no such consistent enrichment, with only a single category showing downregulation (Flagellum, p<9.62E-3). This indicates that loss of CELF6 *in vivo* dysregulates both nuclear transcription and expression of key synaptic genes.

## Discussion

In this study, we have provided the first identification of binding targets of CELF6 *in vivo*. CELF6 primarily binds to 3’UTR regions of mRNAs, similar to CELF4 (31). 3’UTR elements under CELF6 CLIP peaks are enriched for several motifs identified previously by RNACompete (11), including UGU-containing motifs, similar to CELF1 preferences (6). Additionally, using PTRE-Seq (25), we are able to evaluate systematical the impact of CELF6 across hundreds of binding elements. We show that CELF6 and other CELFs generally down-regulate these 3’UTR elements *in vitro*, and this can be abolished by motif mutation. Although CELF6 has been shown to regulate alternative splicing in skeletal muscle (10), we find very few significant binding events outside of 3’UTR regions, suggesting that CELF6 in the brain may be more involved with post-splicing regulation of mRNAs. Additionally, we found few effects to TE in our reporter assays, suggesting that CELF6’s repression of targets is mediated primarily by enhancement of mRNA degradation. Finally, we show these same targets are generally similarly regulated *in vivo*.

Our work here raises a number of interesting biological questions. When transfected *in vitro*, where CELF6 exerts an effect, it is associated with lowered mRNA abundance. However while most of the library was clearly repressive, many elements were not further sensitive to CELF6 expression. This suggests that either CELF6 has additional functions on these transcripts not assessed in our assays (e.g. RNA localization), or that its functional impact on these sequences might depend on the cellular context with regard to the expression of other genes (e.g. other RBPs or miRNAs). While CELF2-6 RNA levels were largely undetectable in SH-SY5Y cells, the complement of other RBPs is likely to differ in the brain and even between different populations of cells in which CELF6 is expressed. We did detect levels of MBNL1 in our SH-SY5Y cells, an RBP which has been shown to antagonize CELF1 (6) leading to mRNA stabilization. Additionally, it has also been shown that with regard to splicing function, CELF1, CELF2, and CELF6 can all antagonize MBNL1 function (32). If MBNL1 and CELF6 can also act antagonistically, then the resistance to CELF6 of some constructs in culture may be due to competition from this or similar proteins. Antagonistic activity of RBPs on mRNA translation and stability has also been observed for CELF2 and HUR (33) and ELAVL1 and ZFP36 (34). Antagonistic activity of RBPs may be mediated by the proximity of binding sites in the 3’UTR, with specific knockdown of RBPs freeing access to other RBP binding sites and resulting in changes to mRNA levels, as has been found for UTRs containing AU-rich elements (35). In our analysis of CELF6 CLIP targets, we found several binding motifs showing enriched abundance in these UTRs for RBPs of different families. Future work with new libraries specifically designed for dissecting the interaction of these motifs may explain why some elements are more sensitive to CELF overexpression than others in these cells.

We found that CELF6 expression was associated with down-regulation of mRNA abundance of library elements. As all reporter elements contain the same promoter, any alterations are unlikely due to changes to transcriptional activity. The simplest explanation is that CELF6 might enhance mRNA decay. Indeed, CELF1 has been shown able to recruit poly A ribonuclease (PARN) to RNA targets to facilitate mRNA decay, and Moraes and colleagues found that CELF1 could associate with PARN *in vitro* (Moraes et al., 2006). Thus the simplest model is that all CELFs can assume similar actions, especially given our finding that CELFs3-6 can all induce repression of the same reporter elements. Additionally, the Xenopus homologue of the CELF proteins, EDEN-BP, has also been shown to regulate deadenylation, and that oligomerization of the protein is required for this activity (36). It is currently unknown whether any of the mammalian CELF proteins are able to hetero or homo oligomerize and the extent to which this may affect the functional activity of these proteins.

It is also interesting to speculate that CELF6 may have roles beyond regulating ribosome occupancy, translation efficiency, and transcript abundance that were assessed here. Indeed, the enrichment of synaptic protein mRNAs targeted by CELF6 hint there could also be a potential role for regulating mRNA localization. Neurons are known to carefully regulate mRNA localization to allow local translation to influence the development and strength of specific synaptic connections. However, as CELF6 expression is limited to fairly sparse populations of neuronal cells, none of which have been extensively studied with regards to local translation, much foundational work would be required before these questions could be readily addressed.

Finally, we have presented the first identification of CELF6 binding targets *in vivo* and shown that CELF6 decreases translation by decreasing mRNA abundance, both *in vitro* and *in vivo*. Some of these targets may mediate the behavioral consequences of CELF6 mutation, which include both communicative and exploratory deficits (8), as well ablation of cocaine mediated reward (Maloney et al, under review). Together, our findings indicate that mRNA translation must be carefully tuned for normal brain function, and that even subtle disruption of mRNA levels can substantially change organismal behavior. The data provided here present an opportunity for further investigations into which of these mRNA targets, in which cell types in the brain, regulate these behaviors.

## Materials & Methods

### Experimental Model and Subject Details

#### Animal Models

All protocols involving animals were approved by the Animal Studies Committee of Washington University in St. Louis. Cages were maintained by our facility on a 12 hr: 12 hr light:dark schedule with food and water supplied ad libidum.

#### Genotyping

Genotyping of all mice was performed using a standard protocol. Animals were genotyped from toe clip tissue lysed by incubation at 50°C in Tail Lysis Buffer for 1 hour to overnight (0.5 M Tris-HCl pH 8.8, 0.25 M EDTA, 0.5% Tween-20) containing 4 *μL*/mL 600 U/mL Proteinase K enzyme (EZ BioResearch), followed by heat denaturation at 99°C for 10 minutes. 1 *μL* Crude lysis buffer was used as template for PCR with 500 nM forward and reverse primers, as specified, using 1X Quickload Taq Mastermix (New England Biolabs) with the following cycling conditions: 94°C 1 min, (94°C 30 s, 60°C 30 s, 68°C 30 sec) x 30 cycles, 68°C 5 minutes, 10°C hold. All referenced PCR primer sequences listed below are found in Supplemental Table S9: Key Resources.

#### Mice for CLIP

Mice used in CLIP experiments derived from 4 litters of CELF6-HA/YFP x C57BL6/J crosses. Each sample used for sequencing was generated by pooling tissue from 3-4 CELF6-HA/YFP+ animals or WT animals from each litter. Pooling tissue was required to generate sufficient material for successful CLIP library generation. Genotyping was performing using HA-F/YFP-R pair (presence or absence of HA-YFP cassette) & Actb-F/Actb-R pair (internal PCR control). Genotyping was performed on post-natal day 7 to select animals for pools, and then performed again on tissue collected after sample processing on post-natal day 9 to confirm genotypes. Sex of animals was noted but not controlled in statistical analysis of CLIP-Seq data as all pools contained animals of both sexes.

1. Litter 1: 8 total animals. <u>CELF6-HA/YFP Pool 1</u>: 4 animals: 1 male & 3 females. <u>CTL Pool 1</u>: 4 animals: 3 males & 1 female.
2. Litter 2: 8 total animals. <u>CELF6-HA/YFP Pool 2</u>: 4 animals: 2 males & 2 females. *CTL Pool from this litter was excluded because upon confirmatory regenotyping it was determined that 1 CTL animal in the pool was actually CELF6-HA/YFP+*.
3. Litter 3: 6 total animals. <u>CELF6-HA/YFP Pool 3</u>: 3 animals: 2 females & 1 male. <u>CTL Pool 2</u>: 3 animals: 2 males & 1 female.
4. Litter 4: 8 total animals. <u>CELF6-HA/YFP Pool 4</u>: 4 animals: 2 females, 2 males. <u>CTL Pool 3</u>: 4 animals: 3 males, 1 female.

#### Mice for Agilent Expression microarray

8 Celf6+/+ (WT) (3 males & 5 females) and 8 Celf6 ^-/-^ (KO) animals (4 males & 4 females) derived from 13 litters of Celf6^+/-^ X Celf6 ^+/-^ crosses were used to generate tissue for the RNA microarray experiment used to assay expression of CELF6 CLIP targets. Animals were genotyped as above using Celf6genoF/Celf6genoR primer pair and age when tissue was harvested ranged between 3.5-9 months.

#### Cell Culture

SH-SY5Y neuroblastoma cells (ATCC CRL-2266) were maintained at 5% CO2, 37°C, 95% relative humidity in 1:1 Dulbecco Modified Eagle Medium/Nutrient Mixture F-12 (DMEM/F12 Gibco) supplemented with 10% fetal bovine serum (FBS Sigma). Under maintenance conditions, cells were also incubated with 1% Penicillin-streptomycin (Thermo), but antibiotics were not used during transient transfections. Cell passage was performed with 0.25% Trypsin-EDTA (Thermo).

### Method Details

#### CLIP

Our CLIP procedure is modeled after the procedure of Wang et al. (6) and from personal communication with the laboratory of Eric Wang. Post-natal day 9 mice were euthanized by rapid decapitation, and brains were dissected. Cortices and cerebella were removed, retaining basal forebrain, striatum, diencephalon, colliculi, and hindbrain regions, which are the brain regions with highest CELF6 expression (9). Dissected tissue was flash frozen in liquid nitrogen and then powdered with a mortar and pestle cooled with liquid nitrogen and kept on dry ice in 10 cm Petri dishes until use. Crosslinking was performed using 3 rounds of 400 mJ/cm^2^ dosage of 254 nm ultraviolet radiation, with petri dishes on dry ice, in a Stratalinker UV crosslinker. After each round of crosslinking, powder in the dishes was redistributed to allow for even crosslinking. After crosslinking, powders were kept on wet ice and incubated with 1mL lysis buffer (50 mM Tris-HCl pH 7.4, 100 mM NaCl, 1X c0mplete EDTA-free protease inhibitor (Sigma), 0.04 U/μL recombinant RNasin (Promega), 10 mM activated sodium orthovanadate, 10 mM NaF). Recombinant RNasin does not inhibit RNase I which was used for RNase digestion in CLIP and was added to prevent other environmental RNase activity. To obtain both cytoplasmic and nuclear fractions in the lysate, lysis buffer was supplemented with NP40 (Sigma CA630) detergent (final concentration 1%) and subjected to mechanical homogenization in a teflon homogenizer 10 times, and lysates were allowed to incubate on ice for 5 minutes. For RNase digestion, RNase I_f_ (New England Biolabs) was diluted to final concentration 0.5, 0.1, or 0.05 U/mL per lysate for radiolabeling. For control radiolabeled sampels (no crosslink and WT tissue immunoprecipitates), the highest (0.5 U/mL) concentration of RNase was used. For samples used for sequencing, 0.05 U/mL final concentration was used. RNase-containing lysates were incubated in a thermomixer set to 1200 RPM at 37°C for 3 minutes and then clarified at 20,000xg for 20 minutes. 2 % input lysate was saved for input samples for sequencing. Per immunoprecipitation, 120 μL of Dynabeads M280 streptavidin (Thermo) were incubated with 17 μL 1 mg/mL biotinylated Protein L (Thermo), and 36 μg each of mouse anti-EGFP clones 19F7 and 19C8 antibodies (MSKCC) for 1 hour. Beads were prepared in batch for all immunoprecipitations and then washed five times with 0.5% IgG-free bovine serum albumin (Jackson Immunoresearch) in 1X PBS, followed by three washes in lysis buffer. Clarified lysates were incubated with coated, washed beads for 2 hours at 4°C with end-over-end rotation and then washed in 1mL of wash buffer (50 mM Tris-HCl pH 7.4, 350 mM NaCl, 1% NP-40, 0.04U/μL RNasin) four times, for 5 minutes with end-over-end rotation at 4°C. For radiolabeling experiments, 60% of washed bead volume was reserved for immunoblotting and added to 20 μL of 1X Bolt-LDS non-reducing sample buffer (Thermo), and 40% proceeded to radioactive labeling. Beads for radioactive labeling were subsequently washed 3x200 μL in PNK wash buffer (20 mM Tris-HCl pH 7.4, 10 mM MgCl2, 0.2% Tween-20, 2.5 U/μL RNasin) and then incubated with 10 μL PNK reaction mixture (1X PNK reaction buffer (New England Biolabs), 4 μCi γ^32^P-ATP (Perkin Elmer), 10 U T4 PNK (New England Biolabs)) for 5 minutes, at 37°C. After labeling, samples were washed in 3x200 μL PNK wash buffer to remove unincorporated label, and then added to 10 μL of 1X Bolt LDS non-reducing sample buffer. All samples in sample buffer were heated for 10 minutes at 70°C and then separated on a 4-12% gradient NuPAGE Bis/Tris gels (Thermo) and then transferred to PVDF membranes with 10% methanol for 6 hours at constant 150 mA. Samples for immunoblot were blocked for 1 hour in block solution (5% nonfat dried milk in 0.5% Tween-20/1X TBS), and then overnight with 1:1000 chicken anti-GFP antibodies (AVES) with rocking at 4°C. Blots were washed 3x5minutes in 0.5% Tween20/1X TBS and then incubated with 1:5000 anti-chicken HRP secondary antibodies (AVES) for 1 hour at room temperature and treated with Biorad Clarity enhanced chemiluminescence reagents for 5 minutes and chemiluminescent data acquired with a Thermo MyECL instrument. Radioactive signal was acquired using an Amersham Typhoon Imaging System and a BAS Storage Phosphor screen (GE Healthcare Life Sciences).

#### CLIP-Seq Sequencing Library Preparation

For CLIP-Seq, EGFP immunoprecipitated WT and HA-YFP+ tissue and 2% input samples were purified from PVDF membranes as follows. Membrane slices were cut with a clean razor according to the diagram in Figure 1A, from unlabeled samples as has been performed in eCLIP (14). PVDF membrane slices were incubated in 1.7 mL microcentrifuge tubes with 200 μL Proteinase K buffer (100 mM Tris-HCl pH 7.4, 50 mM NaCl, 10 mM EDTA, 1% Triton-X 100) containing 40 μL of 800 U/mL Proteinase K (NEB) and incubated in a horizontal shaker at 250 RPM, 37°C for 1 hour. Horizontal shaking reduces the need to cut the membrane into small pieces per sample which is seen in many protocols, and 1% Triton-X 100 in the Proteinase K buffer facilitates increased yield from the membrane by preventing binding of Proteinase K to the membrane. 200 μL of fresh 7M Urea/Proteinase K buffer is then added to slices and tubes are incubated an additional 20 minutes with horizontal shaking at 250 RPM, 37°C. RNA is purified by addition of 400 μL of acid phenol/chloroform/isoamyl alcohol and shaken vigorously for 15 s and allowed to incubate 5 minutes on the bench. RNA samples are centrifuged at 20,000xg for 10 minutes. Aqueous layers are purified using a Zymo-5 RNA Clean & Concentrator column. Output from CLIP’d RNA samples was estimated for total concentration using an Agilent Bioanalyzer and approximately 0.5 ng of RNA was used to prepare next generation sequencing libraries. The full protocol for sequencing library preparation is given in Supplementary Protocol 1. Although this protocol is based on eCLIP, Supplemental Protocol 1 is generalized for any RNA-Seq preparation.

#### Total RNA-Seq of SH-SY5Y cells

For preliminary RNA-Seq of SH-SY5Y cells, 8 replicate subconfluent (approximately 80%) 10 cm dishes (TPP) were harvested by dissociation from the plate by pipetting and centrifugation at 500xg, 5 minutes at room temperature. Pellets were lysed according to TRAP protocol lysis conditions (Supplemental Protocol 2) and RNA was purified using Trizol LS, followed by treatment for 15 minutes at 37°C with DNase I (NEB) and cleanup using Zymo RNA Clean and Concentrator-5 columns (Zymo Research). RNA samples were assessed by Agilent Tapestation with RINe values between 8-10. RNA was fragmented and prepared into libraries for next generation sequencing using Supplemental Protocol 1 as for CLIP-Seq samples.

#### Total RNA-Seq and CLIP-Seq Raw Data Processing

As all total CLIP-Seq and Total RNA-Seq samples were prepared using a unified library preparation procedure, raw data processing used the same set of methods and tools (see Supplemental Table S9) for strand-specific quantification of sequencing reads containing unique molecular identifiers (See Supplemental Protocol 1) to collapse amplification duplicates. Briefly, sequenced paired-end 2x40 on an Illumina Next-Seq. Unique Molecular Identifier (UMI) sequences were extracted from Illumina Read 2, and reads were trimmed for quality using Trimmomatic. Using STAR, remaining reads were aligned to ribosomal RNA, and unalignable reads corresponding to non-rRNA were aligned to the mm10 mouse reference genome and assembled into BAM formatted alignment files. BAM files were annotated with UMI information using the FGBio Java package. PCR duplicates assessed by their UMIs were removed from the BAM files using Picard Tools.

#### Peak Calling and Read Counting

For CLIP-Seq Piranha peak calling analysis, the genome was windowed into 100 bp contiguous windows with 50% overlap using Bedtools. To ensure that called peaks would have the same boundaries for counting read density under a peak for each replicate sample, peaks were first called on a merged BAM file across all YFP-HA+ immunoprecipitated CLIP samples. Piranha p-values for significant peaks were adjusted for multiple testing using Benjamini Hochberg, and all peaks called with False Discovery Rate < 0.1 were kept for further analysis and stored as a Gene Transfer Format (GTF) file. In practice we found that Piranha peaks varied quite widely, with some peaks called with widths on the order of several kilobases despite having a clear local maximum. To more narrowly count reads near peak maxima in a consistent way across all peaks, Piranha peak boundaries were truncated to a width of 100 base pairs around the peak maximum. For the UCSC annotated gene based analysis, UCSC table browser was used to generate GTF files containing all annotated exons from 3’UTR, coding sequence (CDS), 5’UTR, as well as intronic regions. In order to ensure correct mapping of reads to splice sites, the table of intron annotations was allowed to overlap the surrounding exons by 10 bases. In the end, this procedure produced 5 GTF files:

- Piranha peaks (100bp around called maxima from merged BAM file)
- 3’UTR
- CDS
- 5’UTR
- Introns

Each of these GTFs was used as a template for strand-specific feature counting in individual samples using the featureCounts program in the Subread package (Supplemental Table S9). SH-SY5Y Total RNA-seq samples were counted based on a GTF of all UCSC-annotated gene exons to derive total gene counts.

#### CLIP Motif Enrichment Analysis

50 bp regions under the maxima of peaks in CLIP target 3’UTRs were used in MEME Suite (Supplemental Table S9) and compared to 50 bp regions sampled randomly from the 3’UTRs of non-targets - genes which exhibited 0 or negative fold enrichment in YFPHA+ CLIP samples compared to input and WT controls. The DREME tool was used to search *de novo* motifs 6-10 nucleotides in length with E < 0.05. The AME tool was used to search motif enrichment against the CISBP-RNA database. Parameters for AME were: score metric = maximum FIMO score, testing = ranksum test, E<0.05. Peaks were originally called only in the Piranha analysis above. As a number of targets from the annotation-based analysis did not overlap (see Supplemental Table S2 and S3), these 3’UTRs were also scanned for peaks using only the local read density as background.

The position weight matrices (PWMs) for CISBP-RNA motifs showing significant enrichment in CLIP 3’UTR peaks were hierarchically clustered (Figure 2B,2C) by using vectorized PWMs for each motif and the *hclust*() function in R (using the “complete” method). To determine whether or not there were significant matches to a cluster, rather than an individual CISBP-RNA motif PWM, average PWMs were computed by averaging the PWMs across all cluster members. These averaged PWMs were then used with the MEME-Suite FIMO tool to determine whether individual peaks had significant matches to each of the clusters (Supplemental Table S4).

#### PTRE-Seq Reporter Library Preparation

All oligos used for library preparation are shown in the Supplemental Table S9. We generated the pmrPTRE-AAV backbone from an existing mtdTomato construct by PCR amplification and subcloning of the following elements: CMV promoter and a T7 promoter, PCR amplified and subcloned into the MluI restriction site of pQC membrane TdTomato IX. CMV and T7 promters were amplified from pcDNA3.1 using pCMV_T7-F/ pCMV_T7-R and Phusion polymerase (NEB). Then, in order to add a NheI-KpnI restriction enzyme cassette into the 3’UTR, the entire pmrPTRE-AAV plasmid was amplified (pmrPTRE_AAV_Full_F/R) and recircularized using Infusion HD (Clontech). The correct backbone sequence of pmrPTRE-AAV was confirmed by Sanger sequencing.

Originally 473 120 bp sequences under CLIP Peaks were considered for cloning into the library across significant genes. Mutations to motifs found by AME were made as follows. The location of significant matches to motifs were determined using FIMO.

Next, for each matched motif, the PWM representing this motif in CISBP-RNA was used to determine the choice of mutation at each base. Only bases showing a probability of 0.8 or greater were mutated at any position. Bases showing PWM probability >0.8 were mutated to the base showing the minimum value of PWM at that position. If all other three bases showed equal probability, a base was randomly selected from the three. The procedure was repeated at each position for each motif match, before moving to the next matching motif. Where motifs overlapped, lower ranking motifs (based on FIMO score) did not override mutations already made based on higher ranking motifs. After completion this generated a set of 473 mutant elements which ranged between 1 - 25 mutated nucleotides. Subsequently, these sequences were scanned for poly A signals and restriction enzyme sites which would interfere with cloning (NheI, KpnI), which were removed. A final set of 436 120 bp sequences, attached to 6 unique 9 base pair barcodes, as well as priming sites for amplification and cloning (final length 210 bp), and a paired set of 436 mutant sequences similarly prepared were synthesized by Agilent Technologies.

Obtained synthesized sequences were amplified with 4 cycles of PCR using Phusion polymerase with primers GFP-F and GFP-R. We selected these priming sites as these are standard primers in our laboratory used for genotyping that result in robust amplification. The library was PAGE purified and concentration of recovered library was estimated by Agilent TapeStation. The library was digested with NheI and KpnI enzymes and ligated into pmrPTRE-AAV with T4 Ligase (Enzymatics). In order to ensure high likelihood of obtaining all library elements, we prepared our plasmid pool from approximately 40,000 colonies.

#### PTRE-Seq Reporter Library Transfection

His/Xpress-tagged CELF3,4,5,&6 were obtained from the laboratory of Thomas Cooper. For four plasmid experiments, 2500 ng containing equimolar: 2xHis/Xpress-CELF constructs, 1 EGFP-RPL10a, and CELF6 PTRE-Seq library were prepared with Lipofectamine 2000 in Optimem-I (Gibco). For three plasmid experiments, remaining mass was made up with empty pcDNA3.1-His. SH-SY5Y cells wer trypsinized and incubated in 10 cm dishes with Lipofectamine/DNA complexes overnight in DMEM/F12 supplemented with 10% FBS. The following day media was replaced with fresh DMEM/F12 supplemented with 10%FBS and cells were pelleted for Translating Ribosome Affinity Purification (TRAP) and total RNA extraction 40 hours post-transfection. TRAP and total RNA extraction were performed according to the Supplemental Protocol 2 with RNA quality assessed by Agilent TapeStation and all samples had RINe values > 8. The procedure for TRAP in Supplemental Protocol 2 is based on (Heiman et al., 2008) with additional modifications that have been optimized in our laboratory. 5 replicates per condition were generated in batches balanced for all conditions. In each case, replicates were transfected from newly thawed aliquots of cells passaged 1 time before transfection to control for cell passage. Read counts from 1 batch were found to cluster separately from all others after sequencing and data from this batch were excluded. The final data were analyzed from 4 replicates per condition.

#### PTRE-Seq Sequencing Library Preparation

PTRESeq sequencing libraries were prepared by cDNA synthesis using pmrPTRE-AAV antisense oligo (Supplemental Table S9) for library specific priming, and Superscript III Reverse Transcriptase (Thermo) according to the protocol shown in Supplemental Protocol 3. After cDNA synthesis, cDNA libraries were enriched with PCR using Phusion polymerase (Thermo), and pmrPTRE-AAV antisense and sense oligos using 18 cycles. In parallel, plasmid pool DNA was also amplified for sequencing the original plasmid pool. Purified PCR products were digested with NheI and KpnI enzymes and ligated to 4 equimolar staggered adapters to provide sequence diversity for sequencing on the NextSeq. Ligated products were amplified with Illumina primers as in CLIP-Seq library preparation (Supplemental Protocol 1) and subjected to 2x40 paired end next generation sequencing on an Illumina Next-Seq.

#### RT-PCR and qRT-PCR

To confirm overexpression of CELF constructs used in PTRE-Seq experiments, 20 ng of total RNA was converted into cDNA using qScript cDNA synthesis kit (Quantabio, kit employes a mix of both oligo-dT and random hexamer priming) and diluted 4-fold. For RT-PCR, 4 µL of diluted cDNA was used in PCR reactions with 500 nM forward and reverse primers, (Supplemental Table S9, Supplemental Figure S1), using 1X Quickload Taq Mastermix (New England Biolabs) with the following cycling conditions: 94°C 1 min, (94°C 30 s, 60°C 30 s, 68°C 30 sec) x 25 cycles, 68°C 5 minutes, 10°C hold, and then separated by 2% agarose and stained with ethidium bromide. For qRTPCR, 4 µL of diluted cDNA was combined with 500 nM His/Xpress-pcDNA-F and His/Xpress-pcDNA-R primers or HsActb-F/HsAcb-R, and 1X PowerUP SYBR Green Master Mix (Thermo). Each sample/primer combination was run in 4 technical replicates on a Viia7 Real Time PCR System (Thermo) using the following cycling program: 50°C 2 min, 95°C 2 min (95°C 1 s, 60°C 30 sec) x 40 cycles, followed by dissociation step: 95°C 15 s 1.6°/s ramp,60°C 1 min 1.6°/s ramp, 95°C 15 s 0.15°/s ramp. Samples were run alongside no template and no reverse transcription controls to ensure reactions were free of non-target contamination, and dissociation curves were inspected to ensure the absence of non-target amplicons. C_T_ values for each sample were averaged across technical replicates and transformed by first computing the ΔC_T_:= C_T_^His/Xpress^- C_T_^HsActb^, and then computing relative log_2_ expression (“ΔΔC_T_”):= - (ΔC_T_ ^sample^ - ΔC_T_^reference^), where the reference was taken to be the average ΔC_T_ across the CELF6 overexpression condition.

#### PTRE-Seq Reporter Validation

For validation of individual library element reporters (Figure 4), CELF6 CDS was subcloned from His/Xpress-CELF6 into pEGFP-C1 using EcoRI and BamHI restriction sites. Individual Vat1l, Hap1, Peg3, Epm2aip1, and Rab18 reference and mutant sequences were synthesized using Integrated DNA Technologies (IDT) gBlocks Gene Fragments and cloned into pmrPTRE-AAV with NheI and KpnI as above.

50 ng equimolar library element reporters with either pEGFP-C1 (EGFP alone) or EGFP-CELF6 was transiently transfected into SH-SY5Y cells in 96-well plates (TPP). 40 hours post-transfection, media was removed and replaced with warm PBS (1.8 mM KH_2_PO_4_, 10 mM Na_2_HPO_4_, 2.7 mM KCl, 137 mM NaCl, pH 7.4). tdTomato and GFP fluorescence were determined by BioTek Instruments Cytation 5 Cell Imaging Multi-Mode Reader with internal temperature maintained at 37°C. 96-well plates were prepared in 4 replicate batches, where each 96 well plate contained 1 replicate well for all 20 conditions (5 reference, 5 mutant reporters, in both EGFP-CELF6 and EGFP-only transfections). Log_2_ transformed fluorescence intensity measurements were z-score normalized on each plate to account for batch-to-batch differences in transfection efficiency and fluorescence intensity. Data in Figure 4 are shown further normalized to the average value for each reporter in the EGFP-only, reference sequence condition. Example epifluorescent images were obtained using a Leica DMI3000 B microscrope with 20X magnification. Monochromatic images were acquired with QCapture software (QImaging), using gain=1, offset=1, exposure=205 ms for both red and green fluorescent filter sets. 16-bit grayscale images were converted to RGB color and minimally brightness-adjusted for presentation using Adobe Photoshop CS2.

#### Agilent Gene Microarray

Brains from eight wildtype and eight Celf6 mutant mice (see Experimental Model and Subject Details: Animal Models) were extracted, frozen in liquid nitrogen, and crushed into a fine powder, from which RNA was extracted using Qiagen RNEasy columns on a Qiacube robot, following manufacturer’s protocol. RNA was DNase I treated, and RNA quantity and integrity was confirmed using Agilent Bioanalyzer. cDNA was prepared and chemically labeled with Kreatech ULS RNA labeling kit (Kreatech Diagnostics) and Cy5-labeled cDNAs were hybridized to Agilent Mouse v2 4x44K microarrays (G4846A-026655). Hybridization of the labeled cDNAs was done in Agilent 2x gene expression hybridization buffer, Agilent 10x blocking reagent and kreatech Kreablock onto Agilent 4x44K V2 microarrays at 65C for 20 min. Slides were scanned on an Agilent C-class Microarray scanner. Gridding and analysis of images was performed using Agilent Feature Extraction V11.5.1.1.

### Quantification and Statistical Analysis

#### Defining CELF6 CLIP targets by differential expression analysis

Currently there is no standard statistical approach for identification of targets from CLIP data. Methods typically include clustering aligned sequences in individual CLIP RNA samples or replicate averages, with varying probabilistic modeling approaches to assessing signal-to-noise in read density but rarely take into account variance across replicates or differential abundance compared to control samples (6, 37–46). Furthermore, CLIP studies frequently fail to account for differences in the starting abundance of possible target mRNAs, thus reported targets are frequently biased towards highly expressed genes (13). Here, we adopted a strategy of counting reads (as described above) and using standard differential expression analysis tools (edgeR) to make statistical inference on counted features in target immunoprecipitated samples compared to controls, with the hypothesis being that true targets will be enriched in target HA-YFP CLIP samples over both WT controls (representing non-specific pull down) and input samples (accounting for differences in starting abundance of possible target mRNAs).

Samples counted using Subread were imported into R using edgeR (Supplemental Table S9) and normalized for both total library size per sample and feature length as Fragments per Kilobase per Million reads (FPKM). To identify a minimum detection level for subsequent analysis, the relationship between standard deviation across CLIP samples and mean log_2_ FPKM was computed and fitted to a spline. Variability across replicates increases dramatically at poor detection level, and the minimum FPKM required for a feature was set to where the standard deviation decayed to half maximal as a threshold (here determined at 2.5 FPKM). We required that all YFP-HA+ CLIP samples have FPKM > threshold across all YFP-HA+ CLIP samples to be included in analysis. Differential testing was then performed in edgeR against read counts deriving from WT samples or YFP-HA+ input samples. We defined CLIP targets as having positive fold change enrichment in YFP-HA+ CLIP samples compared to both WT CLIP and input samples, with nominal edgeR p-values <0.05 from the edgeR two sample negative binomial exact test. We have proceeded throughout our analysis and PTRE-Seq library generation using targets defined this way in order to maximize exploration of the data. However, we also computed Benjamini-Hochberg adjusted False Discovery Rate for each feature/gene and these are summarized in Supplemental Tables S2 and S3. Our analysis has focused on large groups of features rather than individual target genes and thus we have relied on nominal p-values. However, the reported estimates of FDR can be used to prioritize downstream exploration of biological consequence on individual CLIP targets (Supplemental Table S1 and S2).

#### PTRE-Seq Barcode Counting and Normalization

Barcode counts from sequencing read FASTQ files for each element were determined using a Python script. Read counts were imported into R using edgeR and converted to counts-per-million (CPM) to normalize for differences in library size. Elements showing no counts in the DNA plasmid pool sequencing were removed. Expression was then computed as:

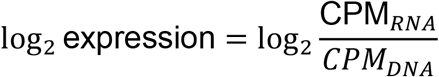

Translation efficiency was computed as:

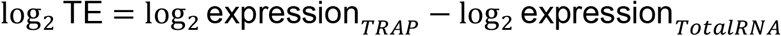

The mean and standard deviation relationship within condition groups were determined for log_2_ expression across elements as in processing for CLIP-Seq in order to filter out poorly detected elements. A minimum log_2_ expression value of -4.29 was determined as a lower bound cutoff which corresponded to approximately 10 counts. We required that all 4 replicates in at least 1 condition had expression levels above this threshold. Finally, after filtering on expression, we required that all elements have at minimum 3 out of the original 6 barcodes present, and present for both reference and mutant alleles.

#### PTRE-Seq Linear Mixed Model Analysis

The final set of elements after filtering on expression and numbers of barcodes was 423 across 172 unique gene UTRs. Individual linear mixed models were performed using lme4 (Supplemental Table S9) fitting the following:

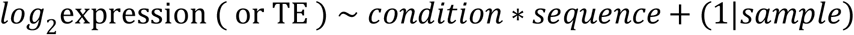

where barcodes were used as repeated measures for each sample, per element. Fixed effect terms of condition referred to either: (a) CTL or CELF6 expression for analyses in Figure 3, or (b) CTL, CELF6, CELF3, CELF4, CELF5, CELF3/6, CELF4/6, CELF5/6 for analyses related to Figure 5. Fixed effect term of sequence was either (a) reference or (b) mutated sequence. Omnibus Analysis of Deviance tests for significant effects of fixed effect terms were computed using likelihood ratio tests in R with the car package (Supplemental Table S9). Estimates of R^2^ were determined according to the procedure of Nakagawa and Schielzeth (47) which provides a simple method for obtaining these estimates from non-linear and empirical model fits in a “percentage of variance explained”-interpretative sense. Omnibus p-values for fixed effects are also reported in Supplemental Tables S6-S8 for models alongside Benjamini Hochberg adjusted False Discovery Rates for TRAP, input expression, and translation efficiency (TE) respectively.

Figures 3 and 5 show analysis of these across all individual models and for subsets of elements with smaller numbers of mutations introduced (≤7 or ≤12) to discern whether radically mutating elements has exerted a strong effect. Because these subsets are nested and because this bird’s-eye view analysis is pooling independently fitted models, we have assessed significance between them using non-parametric Mann Whitney U tests for differences between medians. Post hoc multiple comparisons significance and confidence intervals reported for individual elements in Figure 3 and 4, in order to identify the sources of interaction, were computed using the *multcomp* R package (Supplemental Table S9).

#### PTRE-Seq Validation Analysis

Validation of reporters in Figure 4 was assessed using the car package in R to compute Type II two-way ANOVA with main effects of sequence and CELF6 overexpression and their interaction, with post-hoc pairwise multiple comparisons determined using multcomp.

#### Agilent Expression Microarray Analysis

Features were extracted with Agilent Feature Extraction software and processed with the limma package in R to background correct (using normexp) and quantile normalize between arrays. Genotype of knockout samples was confirmed via decrease of Celf6 mRNA expression level, and differential expression analysis was conducted using limma with empirical bayes (eBayes) analysis. Log_2_ fold change of CLIP targets (defined by the union of both UCSC annotation (3’UTR, 5’UTR’, CDS, introns) and Piranha peak based analysis) was compared to an equivalent number of randomly sampled probes by Welch’s T-test. Gene Ontologies Analysis was conducted in Cytoscape using the Bingo Module, with Benjamini-Hochberg correction and a display cutoff of p<.01. Cellular component results are displayed.

### Availability

All software packages mentioned under Materials and Methods can be found in Supplemental Table S9: Key Resources. UCSC Genome Browser session showing CLIP-Seq data associated with NCBI GEO Accesion GSE118623 (below) can be viewed following this link:

http://genome.ucsc.edu/cgibin/hgTracks?hgS_doOtherUser=submit&hgS_otherUserName=mrieger&hgS_otherUserSessionName=celf6_clip_mm10

### Accession Numbers

Raw and processed sequencing data from CLIP-Seq can be accessed through the National Center for Biotechnology Information’s Gene Expression Omnibus under accession number GSE118623.

## Acknowledgment

We would like to thank Eric Wang, Eric Van Nostrand, Jean Schaffer, and Joshua Langmade for advice on CLIP protocols and sequencing library preparation. We also thank Kyle Cottrell and Sergej Djuranovic for consultation on the preparation of the PTRESeq assays, and Hemangi Chaudhari, Kristina Sakers, Bernard Mulvey for advice. We also thank Allison Lake, Susan Maloney, Affua Akuffo, Jasbir Dalal, and Nilambari Pisat who were involved in generation of mice, data collection, and advice on this project. We thank the Thomas Cooper laboratory for CELF overexpression constructs.

## Author Contributions

MAR, JDD, DK, and BAC designed experiments. MAR performed experiments. MAR and JDD wrote manuscript. MAR, JDD, DK, and BAC revised manuscript.

## Supplemental Information

**Supplemental Figure S1: Expression of His/Xpress-tagged CELF constructs in PTRE-Seq replicates**. CELF6 CLIP UTR element PTRE-Seq library was transiently expressed in neuroblastoma SH-SY5Y along with EGFP-RPL10a and the following constructs: pcDNA3.1 (CTL), (C6), His/Xpress-CELF3 (C3), His/Xpress-CELF4 (C4), His/Xpress-CELF5 (C5). Constructs were transfected singly or in combination with His/Xpress-CELF6, and overexpression was confirmed by RT-PCR with 25 cycles, using primers for CELF1-6, His-Xpress tag, or ACTB as a loading control, separated by 2% agarose, and stained with ethidium bromide. Results shown from (A) Replicate set #1, (B) Replicate set #2, (C) Replicate set #3, (D) Replicate set #4. (E) Quantitative real-time PCR (40 cycles) showing log_2_ expression level of constructs (relative to ACTB), using the His-Xpress tag primer set across conditions, and normalized to the average of the C6 condition. 2 out of 4 CTL samples showed amplification with His/Xpress tag primers in excess of 35 cycles, the remaining 2 samples did not show any amplification. Points show individual sample values and lines show means.

**Supplemental Table S1: Sequencing results for CLIP-Seq samples.** Table showing CLIP-Seq samples 1-4, CLIP-Seq input samples 1-4, and WT CTL IP samples 1-3 with: total read pairs (millions) surviving quality trimming, % duplication as estimated by unique molecular identifiers, uniquely aligning reads (millions), % of uniquely aligning reads aligning to 3’UTR, CDS, 5’UTR, or intronic subgenic regions.

**Supplemental Table S2: Differential analysis of CLIP targets, Piranha peaks.** Table showing the output of edgeR differential expression analysis on read counts deriving from peaks called by Piranha, including: “chr” (chromosome), start, end, strand, gene symbol, geneFeature (utr3, cds, utr5, intron), yfpVinput.logFC (average log_2_ fold change in CLIP samples vs. input), PValue (edgeR negative binomial exact P for CLIP samples vs. input), padj (Benjamini Hochberg adjusted FDR), yfpVwt.logFC (average log_2_ fold change in CLIP samples vs. WT controls).

**Supplemental Table S3: Differential analysis of CLIP targets, UCSC-derived subgenic regions.** Table showing the output of edgeR differential expression analysis as Supplemental Table S2 for read counts deriving from sums of reads in UCSC-annotated 3’UTR, 5’UTR, CDS, and intronic regions of genes.

**Supplemental Table S4: Presence of Celf-, U-rich, and Pcbp3 motif clusters in CELF6 CLIP targets**. For peaks in MEME enrichment analysis, each peak (names “gene_p#” in the case that more than 1 peak was called in the 3’UTR), is listed with the presence (determined by FIMO) “+” or absence “-” of significant matches to Celf-, U-rich-, or Pcbp3-clusters as shown in Figure 2, alongside: “chr” (chromosome), start, and stop locations of the peak, and strand.

**Supplemental Table S5: CELF6 CLIP target sequences used to generate PTRESeq library.** Table showing sequences (“gene_p#” as in Supplemental Table S4) and associated data: “clipFC_yfp_vs_input” (fold change in CLIP samples over input), “utr_name” (gene symbol + “.#” in the event that a gene shows more than one annotated UTR), “gene_name”, chr (chromosome), peakcenter (location of peak maximum), “meme_lower/upper” (start and end coordinates for MEME Suite analysis, 50 bp fragments), “library_lower/upper” (start and end coordinates for PTRE-Seq library generation, 120 bp fragments), strand, reference sequence, mutant sequence, AME output: CISBP-RNA motif ID matches, CISBP-RNA motif alternate names, CISBP-RNA motif alternate “short” names, start/end sequence coordinates of matches, AME adjusted p-value, FIMO scores of matches, FIMO p-values, motif match sequences.

**Supplemental Table S6: PTRE-Seq Linear Mixed Model Results: TRAP.** Linear mixed model results for interaction of sequence (reference, mutant), and CELF6 overexpression condition (CTL, CELF6) using barcode as repeated measure, for each sequence (“gene_p#” as in previous Supplemental Tables S4, S5) in TRAP (ribosomally-association) fraction. “p_sequence” p-value from likelihood ratio test for main effect of sequence, “p_condition” p-value from likelihood ratio test for main effect of condition, “p_interaction” p-value for interaction, “est_r2” the Shinichi-Nakagawa LMM estimate for R^2^, “pctvar_sequence” the estimated percentage of variance explained by main effect of sequence, “pctvar_condition” the estimated percentage of variance explained by condition, “pctvar_interaction” the estimated percentage of variance explained by the interaction term, “pctvar_samples” the estimated percentage of variance explained by the random variance (sample random intercept) term, “pctvar_unexplained” the estimated unexplained variance, “FDR_” the Benamini-Hochberg adjusted FDR for main effects and interaction.

**Supplemental Table S7: PTRE-Seq Linear Mixed Model Results: Input.** LMM results as Supplemental Table S7 for input (total) PTRE-Seq library expression.

**Supplemental Table S8: PTRE-Seq Linear Mixed Model Results: Translation Efficiency.** LMM results as Supplemental Table S7 for PTRE-Seq library translation efficiency.

**Supplemental Table S9: Key Resources.** Table of key resources and links to further information and cited literature, including antibodies, plasmids, oligonucleotides, and software packages.

**Supplemental Protocol S1: Unified CLIP-Seq and RNA-Seq library preparation protocol.** Next generation sequencing library prep protocol with unique molecular identifiers, based on eCLIP, streamlined and generalized for use with any fragmented RNA-Sequencing sample.

**Supplemental Protocol S2: Translating Ribosome Affinity Purification**. Protocol for TRAP as streamlined/optimized in our laboratory.

**Supplemental Protocol S3: PTRE-Seq sequencing library preparation protocol**. Protocol for preparing output of PTRE-Seq library transfections for assay by next generation sequencing.

